# A Mitochondrial Basis for Tead4 Bioavailability at the First Mammalian Cell Fate Decision

**DOI:** 10.64898/2026.03.22.713230

**Authors:** Hannah Sheehan, Andrew J. Piasecki, Dori C. Woods, Jonathan L. Tilly

**Affiliations:** Department of Biology, Northeastern University, Boston, MA; SauveBio, Inc., Worcester, MA

**Keywords:** Tead4, trophectoderm, inner cell mass, mitochondrial heterogeneity, mitochondrial membrane potential, preimplantation embryo, cell fate specification, FAMS

## Abstract

Specification of the inner cell mass (ICM) and trophectoderm (TE) at the first mammalian cell fate decision requires the transcription factor Tead4, yet what restricts Tead4 activity to presumptive TE cells remains unknown. Tead4 localizes to mitochondria, and the ICM and TE harbor distinct mitochondrial populations, but whether Tead4 distribution varies across mitochondrial subtypes in the cleavage-stage embryo has not been examined. Here we used fluorescence-activated mitochondrial sorting (FAMS) to characterize mitochondrial subpopulations in mouse metaphase-II oocytes and 8-cell embryos with respect to size, mitochondrial membrane potential (ΔΨm), and Tead4 protein content. Mitochondria are heterogeneous in size and ΔΨm in both developmental stages, with large mitochondria exhibiting markedly higher ΔΨm than small mitochondria. Tead4 protein is concentrated in the large, high-ΔΨm mitochondrial subpopulation in 8-cell embryos, with 75% of large mitochondria containing Tead4 compared to only 3% of small mitochondria. The overall size distribution of the mitochondrial pool is maintained between oocytes and 8-cell embryos; Tead4 accumulation within the large mitochondrial fraction is therefore a developmentally regulated process initiated specifically during the early embryogenesis. These findings establish for the first time that Tead4 localizes preferentially to large, high-ΔΨm mitochondria in the cleavage-stage embryo, providing a previously unrecognized cellular basis for understanding how Tead4 bioavailability may be regulated prior to TE specification.

## Introduction

Deciphering the mechanisms that drive cell lineage specification has broad implications for understanding pluripotency, stem cell biology, and implantation failure. In mammals, the first cell fate decision occurs at the morula-to-blastocyst transition, when totipotent blastomeres commit to either the inner cell mass (ICM), the source of embryonic stem cells and the future embryo proper, or the trophectoderm (TE), which gives rise to the placenta (Marikawa and Alarcón, 2009; Cockburn and Rossant, 2010; Oron and Ivanova, 2012). Two complementary models have been advanced to explain how this lineage dichotomy is established. In the positional model, outer blastomeres adopt TE identity while inner blastomeres form the ICM based on their relative location within the embryo (Marikawa and Alarcón, 2009). In the polarity model, asymmetric cell division distributes cytoplasmic determinants differentially between apical and basolateral daughter cells, with the apical daughter contributing to the TE and the basolateral daughter to the ICM (Graham and Zernicka-Goetz, 2016; Mihajlovic and Bruce, 2017). The identity of the upstream cytoplasmic determinant(s) driving this asymmetry remains unknown.

Among the earliest molecular markers of TE identity is TEA domain transcription factor 4 (Tead4), expressed from the 4-cell stage onward and required for activation of key TE-specification genes including caudal-type homeobox 2 (Cdx2) and Eomesodermin (Eomes) (Marikawa and Alarcón, 2009; Cockburn and Rossant, 2010; Oron and Ivanova, 2012). Embryos lacking Tead4 fail to activate Cdx2, do not form TE, and are not viable (Yagi et al., 2007; Nishioka et al., 2008). Tead4 activity is modulated by the Hippo signaling pathway: in inner blastomeres, biomechanical strain from cell-cell contacts drives Lats1/2-mediated phosphorylation and cytoplasmic sequestration of the Tead4 co-activator Yes-associated protein (YAP), whereas outer blastomeres with cell-contact-free apical surfaces have reduced Hippo activity, permitting nuclear YAP-Tead4 interaction and TE gene activation (Nishioka et al., 2009; Sharma et al., 2021). However, cell-contact-free apical surfaces exist throughout cleavage, yet blastomere fate only becomes restricted at the 32-cell stage (Marikawa and Alarcón, 2009; Graham and Zernicka-Goetz, 2016; Mihajlovic and Bruce, 2017). Furthermore, the absolute level of Tead4 protein governs its subcellular localization, with enforced overexpression driving nuclear Tead4 and aberrant TE specification even in inner blastomeres (Home et al., 2012). These observations raise the possibility that Tead4 bioavailability is regulated prior to the 32-cell stage by sequestration in a subcellular compartment from which it is released at a critical threshold.

An intriguing candidate compartment for Tead4 sequestration is the mitochondrion. Tead4 localizes to mitochondria in blastocysts, where it promotes mitochondrial gene transcription and supports blastocoel formation (Kaneko and DePamphilis, 2013; Kumar et al., 2018). The N-terminal disordered region of Tead4 (amino acids 1–42) is composed largely of highly polar residues with structural features associated with mitochondrial targeting and uptake (Wattenberg et al., 2007; Kim and Han, 2018). Mitochondria in mammalian oocytes and preimplantation embryos are heterogeneous in size, ultrastructure, and ΔΨm (Diaz et al., 1999; Van Blerkom et al., 2002), and this heterogeneity is not random. The significance of mitochondrial ΔΨm heterogeneity for cell fate has been established in other stem cell contexts: in hematopoietic stem cells, ΔΨm predicts self-renewal versus differentiation potential (Vannini et al., 2016), and dynamic changes in mitochondrial activity during embryonic hematopoiesis drive lineage-biased progenitor output (Prakash and Inamdar, 2025). Whether analogous principles operate in the earliest cell fate decision of mammalian development has not been investigated. Studies of human and mouse blastocysts have established that cells of the TE contain predominantly large, structurally complex, high-ΔΨm mitochondria, while cells of the ICM harbor mostly small, immature, low-ΔΨm mitochondria (Hashimoto et al., 2017; Van Blerkom, 2004). The origin of this lineage-specific mitochondrial distribution, and whether it reflects a cause or a consequence of lineage specification, has not been established.

These observations converge on an important but unaddressed question: does Tead4 exhibit subpopulation-specific localization within mitochondria of the cleavage-stage embryo, and if so, within which mitochondrial subtypes does it reside? Answering this is a necessary first step toward understanding whether mitochondrial heterogeneity contributes to the regulation of Tead4 bioavailability prior to TE specification. To address this, we employed fluorescence-activated mitochondrial sorting (FAMS) (MacDonald et al., 2019), a nanoscale multi-parametric flow cytometric platform capable of identifying, quantifying, and isolating mitochondrial subpopulations based on size, ΔΨm, and protein content at single-organelle resolution. We characterized mitochondrial subpopulations in mouse metaphase-II oocytes and 8-cell embryos with respect to size, ΔΨm, and Tead4 protein localization. Our findings establish that large mitochondria are predominantly high- ΔΨm in both stages, and that Tead4 is concentrated within this subpopulation in the cleavage-stage embryo, providing the first cellular basis for understanding how mitochondrial heterogeneity may regulate Tead4 bioavailability during preimplantation development.

## Materials and Methods

### Animals and Oocyte/Embryo Collection

All animal procedures were performed in accordance with institutional guidelines. Metaphase-II oocytes and embryos were collected from superovulated C57BL/6 female mice (The Jackson Laboratory, Bar Harbor, ME). Superovulation was induced by sequential intraperitoneal injection of pregnant mare serum gonadotropin (PMSG; 5 IU) followed 48 hours later by human chorionic gonadotropin (hCG; 5 IU). Oocytes were collected from the oviducts 13-14 hours after hCG injection by puncturing the ampulla. 8-cell embryos were generated by in-vitro fertilization (IVF) using established protocols (Morita et al., 2000). Briefly, cumulus-oocyte complexes were inseminated with capacitated sperm from C57BL/6 males, and resultant embryos were cultured in KSOM medium + 0.4% BSA at 37°C in 5% CO_2_ to the 8-cell stage.

### Fluorescence-Activated Mitochondrial Sorting (FAMS)

Mitochondria were isolated and analyzed by FAMS as described in MacDonald et al. (2019). Briefly, oocytes or embryos were disrupted by gentle mechanical homogenization in ice-cold isolation buffer, and the resulting homogenate was subjected to low-speed centrifugation to remove cellular debris, followed by collection of the mitochondria-enriched fraction. Mitochondrial size distribution was determined by forward scatter (FSC PMT) and side scatter (SSC) analysis, with size gates defined as small (≤0.30 μm), intermediate (0.31–0.60 μm), and large (>0.60 μm), consistent with published electron microscopy data for mouse oocyte mitochondria (Piko and Matsumoto, 1976) and proportional to the ultrastructurally defined size categories observed in human preimplantation embryos (Hashimoto et al., 2017).

### Assessment of Mitochondrial Membrane Potential

Mitochondrial membrane potential (ΔΨm) was assessed in metaphase-II oocytes and 8-cell embryos using JC-1 (Thermo Fisher Scientific), a dual-emission fluorescent dye that accumulates in mitochondria and undergoes a fluorescence shift from green (FITC channel, low-ΔΨm) to red/orange (PE channel, high-ΔΨm) as membrane potential increases, as previously described (MacDonald et al., 2019). ΔΨm status within each size-based subpopulation was quantified as the proportion of mitochondria classified as ΔΨm+ based on PE-A fluorescence intensity. Gating strategies and controls were as described in MacDonald et al. (2019).

### Immunoanalysis of Tead4 in Mitochondrial Subpopulations

Tead4 protein was detected in mitochondrial subpopulations isolated from 8-cell embryos by FAMS using a mouse monoclonal antibody against Tead4 (Abcam). This antibody has been rigorously validated for Tead4 immunolocalization in mouse embryos using Tead4-mutant embryos as negative controls and has been used in multiple published mitochondrial localization studies (Home et al., 2012; Rivron et al., 2018; Menchero et al., 2019). Its utility for FAMS-based protein detection was validated in the present study. Tead4 immunoreactivity was detected using a goat anti-mouse IgG Alexa Fluor 594-conjugated secondary antibody (Abcam). The proportion of Tead4-positive mitochondria within each size-based subpopulation was quantified as a percentage of total mitochondria in that fraction.

### Statistical Analysis

Data are presented as mean ± SEM. All FAMS experiments were performed in a minimum of n = 3 independent biological replicates. Statistical comparisons were made using one-way ANOVA with appropriate post-hoc testing; p < 0.05 was considered significant.

## Results

### Mitochondria in mouse metaphase-II oocytes exhibit heterogeneity in size and ΔΨm

FAMS analysis of metaphase-II oocytes confirmed that mitochondria span a size range of approximately 0.2 μm to 0.8 μm (Figure 1A), consistent with prior electron microscopy estimates (Piko and Matsumoto, 1976). The small (≤0.30 μm) and intermediate (0.31–0.60 μm) size fractions together accounted for approximately 85% of total mitochondria, with the large (>0.60 μm) fraction representing the remaining ∼15% (Figure 1B).

**Figure 1.**
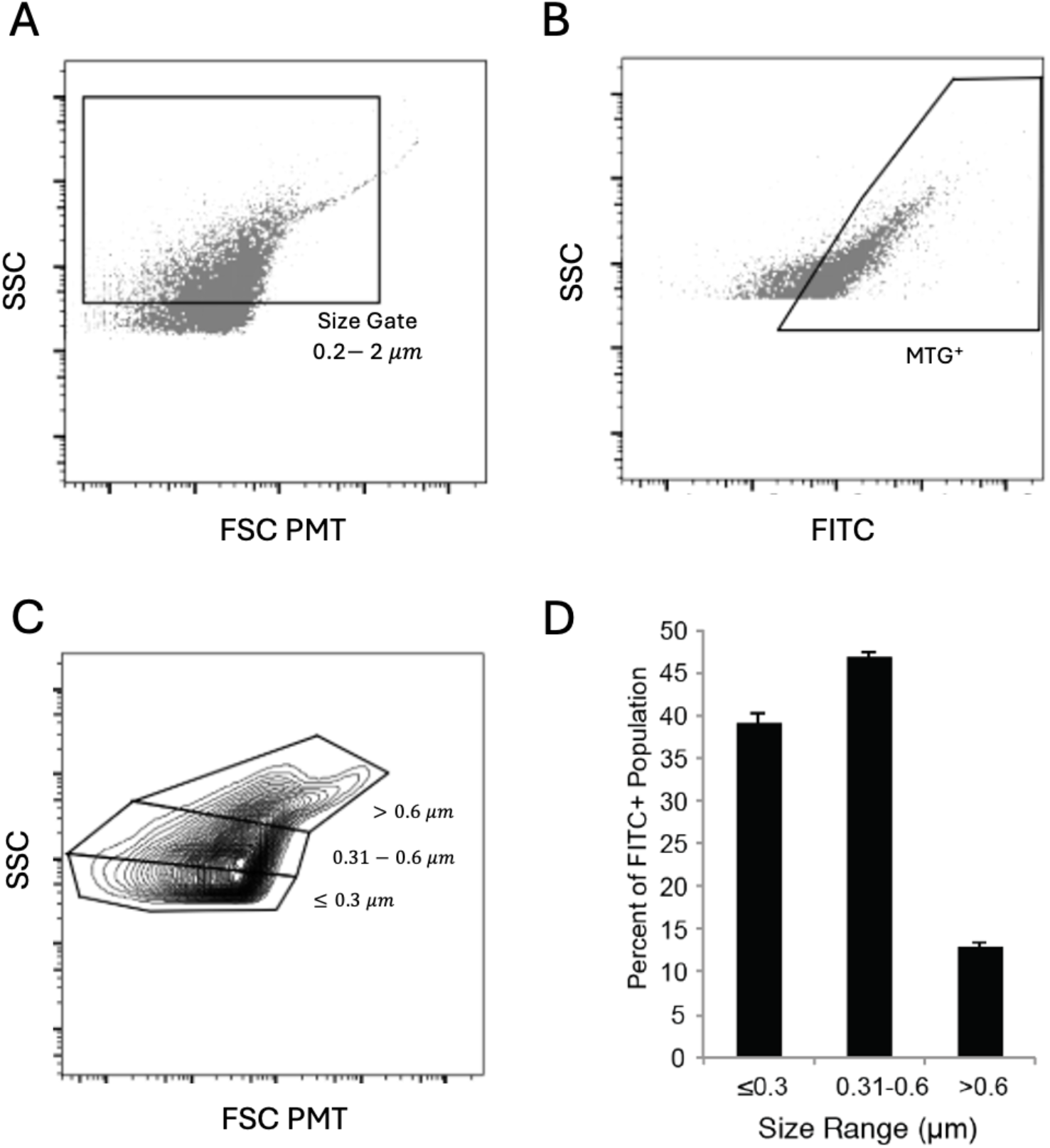
Mitochondrial are heterogeneous in size. Mitochondria were isolated from metaphase-II oocytes and analyzed by fluorescence-activated mitochondrial sorting (FAMS) as described in Materials and Methods. (A) FSC PMT/SSC dot-plot of total events showing the size gate (0.22–2 μm) used to identify the mitochondrial population. (B) FSC PMT/FITC dot-plot confirming that events within the size gate are MitoTracker Green FM-positive, verifying their identity as mitochondria. (C) Representative FSC PMT/SSC contour plot of the MitoTracker Green FM-positive population showing the three size-based subpopulations: small (≤0.30 μm), intermediate (0.31–0.60 μm), and large (>0.60 μm), with the percent of the total mitochondrial population represented by each size fraction indicated. (D) Size breakdown of the mitochondrial population expressed as the percent of the total mitochondrial population represented by each size fraction. Data are presented as mean ± SEM; n = 3 independent biological replicates.

### Large mitochondria exhibit higher ΔΨm than small mitochondria in both oocytes and 8-cell embryos

FAMS-based analysis using JC-1 demonstrated that mitochondria exhibit heterogeneity in ΔΨm in oocytes and 8-cell embryos, with distinct ΔΨm– and ΔΨm+ populations identifiable in each (Figure 2). ΔΨm was not uniformly distributed across size fractions in either stage. In oocytes, 57% of large mitochondria were ΔΨm+, compared with lower proportions in the small and intermediate fractions (Figure 2B). The same size-dependent relationship was observed in 8-cell embryos, where large mitochondria again exhibited the highest proportion of ΔΨm+ mitochondria of all three size fractions (Figure 2D). In both stages, ΔΨm increased progressively with mitochondrial size, with levels in large ΔΨm+ mitochondria exceeding those of small ΔΨm+ mitochondria by more than one log-order, consistent with the greater cristae density of larger mitochondria (Hashimoto et al., 2017; Van Blerkom, 2004; Wolf et al., 2019). The positive relationship between mitochondrial size and ΔΨm is therefore a consistent feature of the mitochondrial landscape from the oocyte through the cleavage stage.

**Figure 2.**
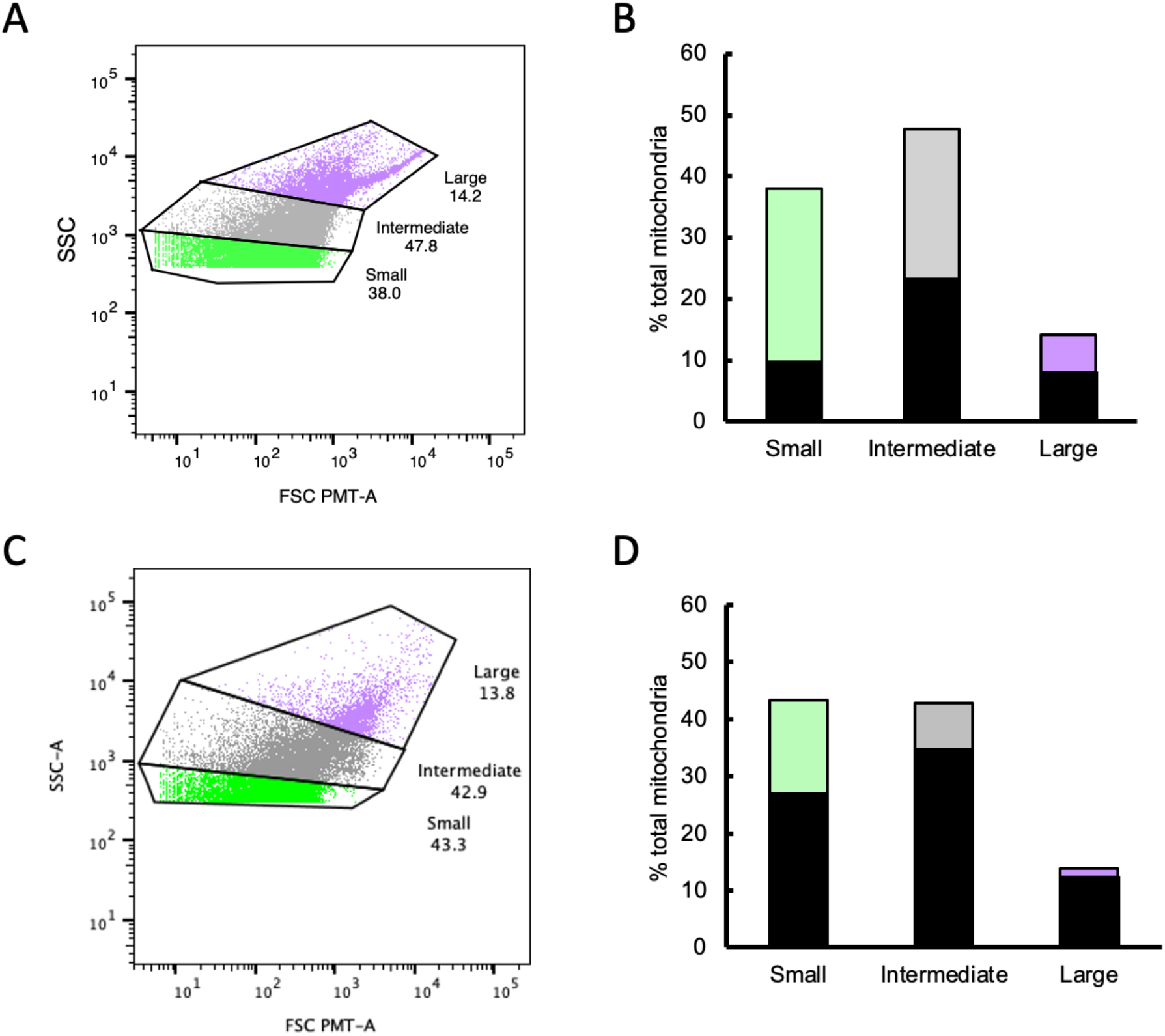
Mitochondrial size predicts functional state. The relationship between mitochondrial size and mitochondrial membrane potential (ΔΨm) was assessed in metaphase-II oocytes and 8-cell embryos by FAMS using JC-1, a dual-emission fluorescent dye that shifts from green fluorescence in low-ΔΨm (ΔΨm–) mitochondria to red/orange fluorescence in high-ΔΨm (ΔΨm+) mitochondria. (A) Representative FSC PMT-A/SSC-A dot-plot of oocyte mitochondria showing the three size-based subpopulations (small, ≤0.30 μm; intermediate, 0.31–0.60 μm; large, >0.60 μm). (B) Proportion of ΔΨm+ (black shading) versus ΔΨm– (colored shading) mitochondria within each size fraction in oocytes. (C) Representative FSC PMT-A/SSC-A dot-plot of 8-cell embryo mitochondria showing the same three size-based subpopulations. (D) Proportion of ΔΨm+ versus ΔΨm– mitochondria within each size fraction in 8-cell embryos. Data are presented as mean ± SEM; n = 3 independent biological replicates per stage. See MacDonald et al. (2019) for gating strategies and controls.

### Tead4 is preferentially localized to large, high-ΔΨm mitochondria in 8-cell embryos

The central objective of this study was to determine whether Tead4 exhibits subpopulation-specific localization within mitochondria of cleavage-stage embryos. We selected 8-cell embryos because Tead4 expression is activated at the 4-cell stage (Nishioka et al., 2008) and Tead4 protein was not readily detected in mitochondrial subpopulations of metaphase-II oocytes by FAMS, consistent with the known transcriptional onset of this gene. At the 8-cell stage, Tead4 is actively expressed yet blastomere fate has not become irreversibly restricted, making this the earliest stage at which mitochondrial Tead4 content can be meaningfully assessed.

FAMS analysis confirmed the presence of small, intermediate, and large mitochondrial subpopulations in 8-cell embryos (Figure 3A), representing approximately 40%, 37%, and 23% of total mitochondria, respectively (Figure 3B). Immunoanalysis of Tead4 across these fractions produced a clear result: 75% of large mitochondria were Tead4-positive, compared to only 3% of small mitochondria (Figures 3B and 3C). Intermediate-sized mitochondria showed intermediate Tead4 immunoreactivity, consistent with a progressive relationship between size and Tead4 content (Figure 3C). Since large mitochondria are predominantly high-ΔΨm in 8-cell embryos (Figure 2C, D), these data establish directly that Tead4 is concentrated within the large, high-ΔΨm mitochondrial subpopulation.

**Figure 3.**
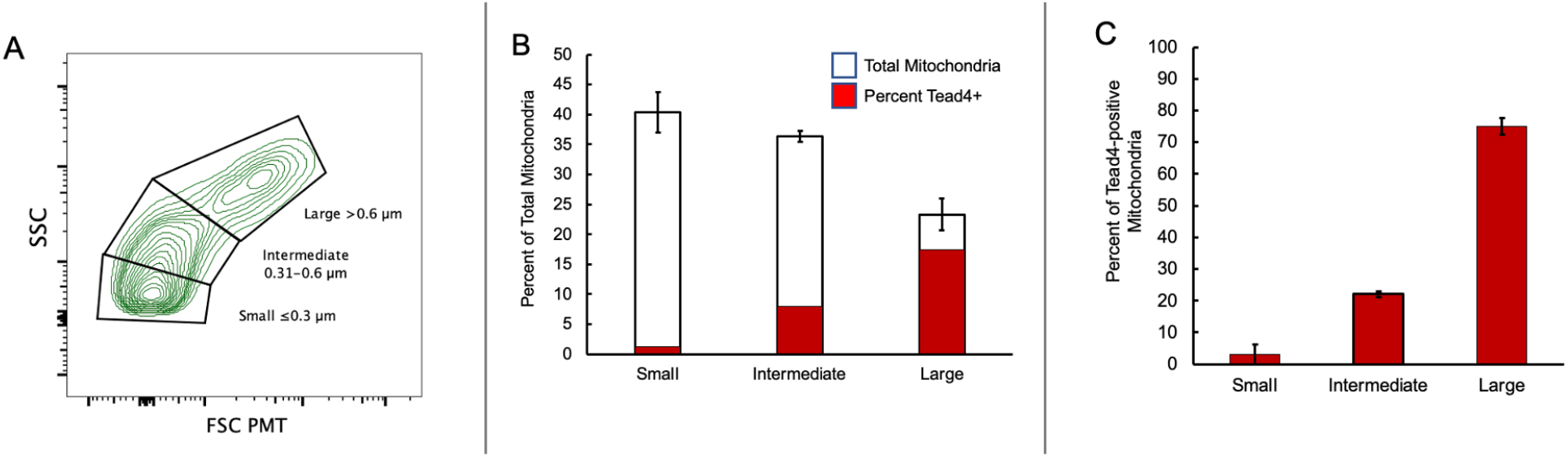
Functional state links to Tead4 localization. Tead4 protein distribution across mitochondrial size subpopulations was assessed in 8-cell embryos by FAMS-based immunoanalysis using a validated monoclonal antibody against Tead4 (Abcam) and an Alexa Fluor 594-conjugated anti-mouse secondary antibody (Abcam). (A) Representative FSC PMT/SSC contour plot showing the three mitochondrial size subpopulations (small, ≤0.30 μm; intermediate, 0.31–0.60 μm; large, >0.60 μm) in 8-cell embryos. (B) Percent of total mitochondria represented by each size fraction (white bars) and the proportion of mitochondria within each size fraction that are Tead4-positive (red bars). (C) Proportion of Tead4-positive mitochondria within each size fraction shown independently. Data are presented as mean ± SEM; n = 3 independent biological replicates. See MacDonald et al. (2019) for methodological details.

### Mitochondrial size distribution is maintained across the oocyte-to-embryo transition and is accompanied by progressive Tead4 accumulation

Comparison of mitochondrial size distributions between oocytes (Figures 1B and 2A, B) and 8-cell embryos (Figure 2C, D) revealed broadly similar proportions of small, intermediate, and large mitochondria across the two stages, indicating that the overall composition of the mitochondrial pool is maintained across the oocyte-to-embryo transition. Despite this similarity in size distribution, the proportion of Tead4-positive mitochondria increases progressively with mitochondrial size in 8-cell embryos (Figure 3C). Since Tead4 expression is not activated until the 4-cell stage (Nishioka et al., 2008), this accumulation of Tead4 within large mitochondria is a developmentally regulated process that occurs specifically during the early embryogenesis.

## Discussion

The primary objective of this study was to determine whether Tead4 exhibits subpopulation-specific localization within mitochondria of the cleavage-stage mouse embryo. This question had not been addressed despite the known mitochondrial localization of Tead4 (Kaneko and DePamphilis, 2013; Kumar et al., 2018) and the well-established heterogeneity of mitochondrial populations in oocytes and preimplantation embryos (Diaz et al., 1999; Van Blerkom et al., 2002; Hashimoto et al., 2017; Van Blerkom, 2004). Tead4 is concentrated in large, high-ΔΨm mitochondria in 8-cell embryos, with 75% of large mitochondria containing Tead4 compared to only 3% of small mitochondria. Large mitochondria exhibit markedly higher ΔΨm than small mitochondria in both oocytes and 8-cell embryos, differing by more than one log-order, and the overall mitochondrial size distribution is preserved across the oocyte-to-embryo transition, with Tead4 accumulating progressively within the large mitochondrial fraction as its expression increases. Together, these findings establish the mitochondrial size- and ΔΨm-dependent distribution of Tead4 as a previously unrecognized feature of the cleavage-stage embryo.

The significance of these findings is best understood in the context of what is known about Tead4 regulation. The absolute level of Tead4 protein governs its subcellular localization, cytoplasmic versus nuclear, and thus its transcriptional activity (Home et al., 2012). Enforced overexpression of Tead4 drives nuclear accumulation and aberrant TE specification even in inner blastomeres that would normally give rise to the ICM (Home et al., 2012), demonstrating that cytoplasmic Tead4 availability is a threshold that must be breached for TE gene activation. Our finding that 75% of large mitochondria in 8-cell embryos contain Tead4 establishes that a substantial depot of this transcription factor is housed within a specific mitochondrial subpopulation well before the 32-cell stage when TE fate becomes fixed. This provides a cellular mechanism by which Tead4 could be sequestered and its bioavailability regulated during the cleavage stage, a possibility that prior studies could not have addressed without subpopulation-level resolution.

The structural attributes of the Tead4 N-terminus are consistent with the preferential mitochondrial uptake we observe. The disordered, highly polar N-terminal region (amino acids 1–42) contains features including transient secondary structure and compositional polarity bias known to facilitate mitochondrial targeting, binding, and import (Wattenberg et al., 2007; Kim and Han, 2018). An established ΔΨm is required for efficient mitochondrial protein import (Zorova et al., 2018), providing a mechanistic basis for why Tead4 accumulates preferentially in large, high-ΔΨm mitochondria as Tead4 expression increases from the 4-cell stage onward (Nishioka et al., 2008). The progressive increase in mitochondrial Tead4 content with increasing organelle size (Figure 3C) is consistent with this, and suggests that Tead4 import is coupled to mitochondrial maturation during the cleavage stage.

The data are consistent with a model in which Tead4-enriched large mitochondria are positioned for asymmetric segregation into outer blastomeres during cleavage. Prior work has shown that subcellular ΔΨm heterogeneity within blastomeres correlates with cell-cell contact geometry: low-ΔΨm mitochondria cluster at intercellular contact regions on the inward-facing surfaces of blastomeres, while high-ΔΨm mitochondria concentrate at cell-contact-free outward-facing surfaces (Van Blerkom et al., 2002). Large Tead4-containing mitochondria, by virtue of their high ΔΨm, would therefore be expected to localize preferentially to apical surfaces of blastomeres, positioning them for preferential segregation into outer daughter cells at each division. Over successive divisions, this would progressively concentrate Tead4 in outermost blastomeres destined to form the TE. Whether this segregation occurs, and whether it is causally linked to TE specification, are questions currently under investigation.

The broader significance of these findings extends beyond the preimplantation embryo. Mitochondrial ΔΨm has emerged as a determinant of stem cell fate in multiple systems: in hematopoietic stem cells, ΔΨm predicts long-term reconstitution capacity and daughter cell fate (Vannini et al., 2016), and dynamic changes in mitochondrial activity during embryonic hematopoiesis drive lineage-biased progenitor output (Prakash and Inamdar, 2025). The present findings extend this principle to the earliest cell fate decision in mammalian development and introduce the subpopulation-specific sequestration of a lineage-specifying transcription factor within mitochondria of defined ΔΨm and size, a feature not previously described in any stem cell context. Mitochondrial subpopulations differing in proteomic profiles, bioenergetic properties, and cell-specific functions (Mootha et al., 2003; Taylor et al., 2003) may represent a broadly conserved mechanism by which mitochondrial heterogeneity shapes stem and progenitor cell identity.

Several questions follow from these findings and are currently under investigation. Whether the localization of Tead4 to large, high-ΔΨm mitochondria is established progressively as Tead4 expression increases from the 4-cell stage, and how Tead4 distribution evolves through successive cleavage stages, remain to be determined. Direct demonstration that large, Tead4-enriched mitochondria segregate preferentially into outer blastomeres, and that this correlates with nuclear Tead4 accumulation and TE gene activation, is the critical next step toward establishing causality. Identifying the trigger responsible for Tead4 release from mitochondria at the 32-cell stage, and testing whether this release drives TE specification, will be necessary to define the full mechanistic pathway.

Tead4 is concentrated in large, high-ΔΨm mitochondria in 8-cell embryos at a ratio of 75% to 3% relative to small mitochondria, a distribution coupled to post-fertilization mitochondrial maturation. These findings establish a previously unrecognized cellular basis for the regulation of Tead4 bioavailability during preimplantation development and open a new line of investigation into the role of mitochondrial heterogeneity in the first mammalian cell fate decision. A conceptual summary of the proposed model is presented in Figure 4.

**Figure 4.**
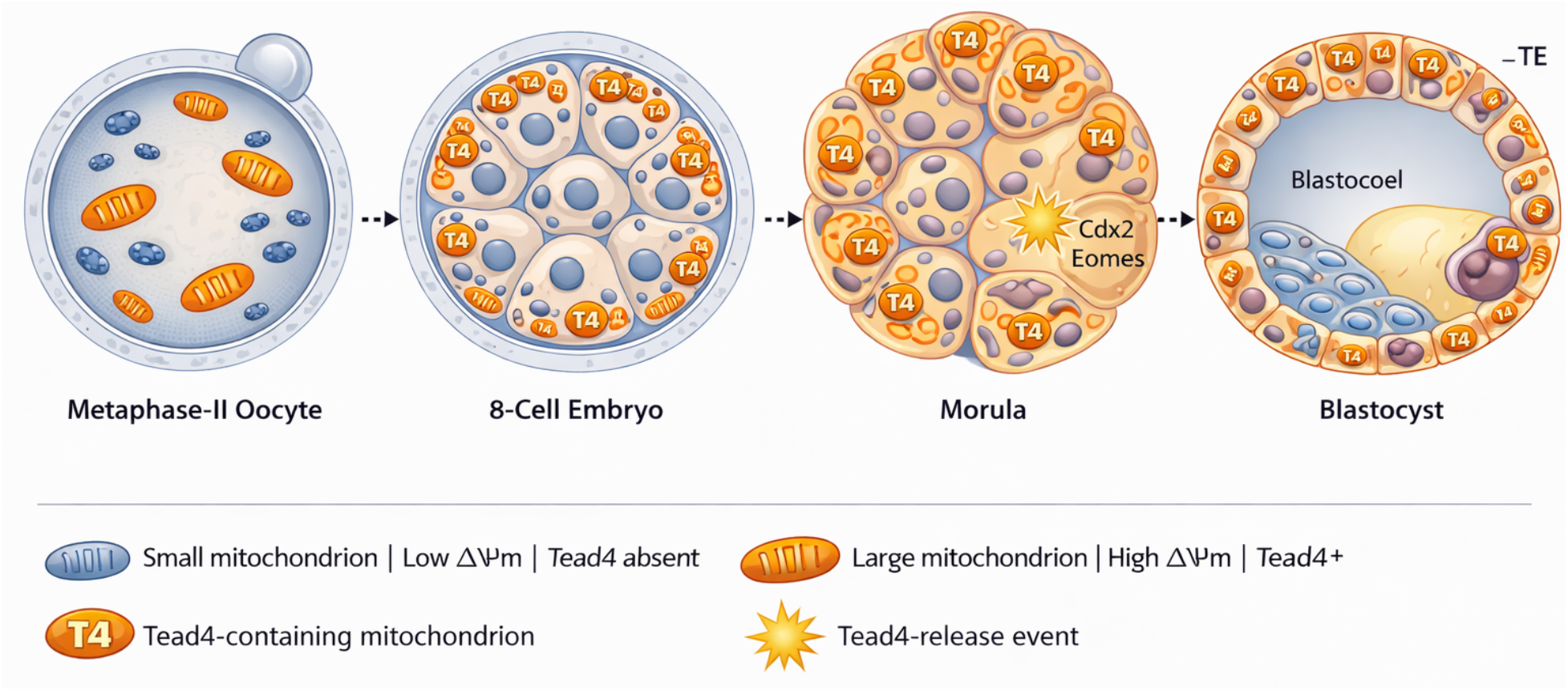
Model for mitochondrial subtype-dependent Tead4 distribution and trophectoderm specification during mouse preimplantation development. Schematic depicting the proposed role of mitochondrial heterogeneity in Tead4 bioavailability at the first mammalian cell fate decision. In the metaphase-II oocyte (left), mitochondria are heterogeneous in size and ΔΨm but Tead4 is not yet expressed. As embryogenesis proceeds through the cleavage stage, shown here at the 8-cell embryo, Tead4 is imported into large, high-ΔΨm mitochondria. The subcellular distribution of ΔΨm, with high-ΔΨm mitochondria concentrated at cell-contact-free apical surfaces and low-ΔΨm mitochondria clustering at intercellular contact zones, positions large Tead4-containing mitochondria for asymmetric segregation into outer blastomeres across successive cleavage divisions. At the morula stage, a threshold event in outer cells triggers release of Tead4 from mitochondria into the cytoplasm, enabling nuclear translocation and activation of TE-specification genes Cdx2 and Eomes. By the blastocyst stage, cells of the trophectoderm (TE) are enriched for large, high-ΔΨm, Tead4-containing mitochondria, while cells of the inner cell mass (ICM) contain predominantly small, low-ΔΨm mitochondria devoid of Tead4. Small blue-gray ovals, small mitochondria with low ΔΨm and no Tead4; large orange ovals with parallel internal cristae, large mitochondria with high ΔΨm and Tead4 enrichment; T4 badge, Tead4-containing mitochondrion; gold starburst, Tead4-release event. TE, trophectoderm; ICM, inner cell mass.

## Acknowledgments

The authors thank Jessica Martin for contributions to data collection and analysis during the early stages of this work. This work was supported by National Science Foundation grant 2227756.

## Author Contributions

Hannah Sheehan performed experiments, analyzed data, and contributed to writing and editing of the manuscript. Andrew J. Piasecki performed experiments. Dori C. Woods and Jonathan L. Tilly conceived and designed the study, supervised all aspects of the work, and wrote the manuscript. All authors reviewed and approved the final version.

